# Plk3 Regulates Bacteremia and Supports Sepsis

**DOI:** 10.1101/2024.02.19.581105

**Authors:** John C Kostyak, Sharath S Sarojini, Meghna U Naik, Wei Dai, James V Michael, Steven E McKenzie, Ulhas P Naik

## Abstract

**Objective:** Sepsis, which is the body’s response to overwhelming infection, can lead to septic shock, characterized by thrombocytopenia, hypotension, and organ damage. Polo-like kinase 3 (Plk3) is a ubiquitously expressed serine/threonine kinase, but its exact role in immune function is unknown.

**Approach and Results:** We used *Plk3^−/−^* and WT mice to evaluate the function of Plk3 in several models of severe sepsis. We found that WT mice die within 48 hours of 100% cecal ligation and puncture (CLP), while *Plk3^−/−^* mice survive. Survival following cecal slurry (CS) injection mirrored that of CLP as recipient WT mice succumbed, while recipient *Plk3^−/−^* mice survived. Analysis of bacterial load 24 hours after CLP revealed that WT blood and peritonea were loaded with bacteria, but bacteria were virtually undetectable in the peritonea or blood of *Plk3^−/−^* mice. To determine if bacteria infiltrate the blood of *Plk3^−/−^* mice shortly after infection, we measured bacteria 1 and 3 hours after CS injection. We found a time-dependent increase in bacteria in the blood of WT mice that was not observed in *Plk3^−/−^* mice. To determine if the lack of bacteria in the blood of *Plk3^−/−^* mice is due to enhanced clearance, we injected *E. coli* IV into WT and *Plk3^−/−^* mice. We found 75% mortality for both WT and *Plk3^−/−^* mice within 72 hours following IV injection suggesting that survival of *Plk3^−/−^* mice following enteric infection is likely due to reduced bacteremia.

**Conclusion:** Collectively our data suggest that Plk3 supports the systemic dissemination of bacteria and subsequent sepsis following enteric infection.

## Introduction

Sepsis is defined as a dysregulated host response to infection that results in organ dysfunction and can lead to septic shock and death. It is the most expensive condition in US hospitals, costing more than $20 billion annually ^1^. In the US 1/3 of all patients that die while in the hospital are septic. Therefore, there remains great need for a better understanding of sepsis and the progression to septic shock. Furthermore, there is currently no therapy available to combat the host response to infection that leads to sepsis.

When local infection is not contained and progresses to systemic infection, a rapid inflammatory response occurs. Sepsis occurs when pathogen associated molecular patterns (PAMPs) such as lipopolysaccharide (LPS) from gram-negative bacteria or lipotechoic acid from gram-positive bacteria interact with pattern recognition receptors (PRRs) such as toll-like receptor 4 (TLR4) and TLR2 on inflammatory cells, respectively ^2^. This causes the release of pro-inflammatory cytokines like tumor necrosis factor-α and interferon-γ, which triggers the release of other pro-inflammatory cytokines like interleukins (IL-1, IL-2, IL-6, IL-8) while suppressing anti-inflammatory cytokines like IL-10. This inflammatory cell response results in a feed forward mechanism that causes super physiological levels of circulating cytokines and triggers the perpetual activation of platelets and inflammatory cells ^3^. Enhanced activation of cells like B-cells and T-cells results in apoptosis and loss of immune cells which contributes to sepsis-induced immunosuppression. It has been shown that circulating lymphocyte counts fall during the onset of sepsis and can remain low in patients for several days. In fact, patients with lymphopenia that persist past the fourth day after sepsis diagnosis are more likely to die than patients that are no longer lymphopenic 4 days after diagnosis ^4^.

Polo-like kinases are a family of five (Plk1, Plk2, Plk3, Plk4, and Plk5) evolutionarily conserved serine/threonine protein kinases. All Plk family members have an N-terminal catalytic domain and two or more C-terminal polo boxes that are responsible for substrate binding ^5^. Plk3, formerly known as Fnk (FGF-inducible kinase), has a cell cycle regulatory function that is dispensable as *Plk3^−/−^* mice are viable and fertile ^6^. However, *Plk3^−/−^* mice are heavier than WT mice and have an increased propensity for tumor development at advanced age ^6^. Several groups provide evidence that Plk3 regulates the cellular stress response to DNA damage, Golgi fragmentation, osmotic stress, and hypoxia ^7–10^. Interestingly, it appears as though, similar to its cell cycle regulatory role, the function of Plk3 in cell stress response may be dispensable ^11^.

Plk3 mRNA and protein expression is induced by adhesion of human monocytes to a cell culture dish and overexpression of Plk3 disrupted the F-actin cytoskeleton ^12^. Plk3-GFP transfection of COS cells shows Plk3 expression at the cell periphery during attachment ^12^. These data are consistent with more recent data that demonstrates a rapid rise in Plk3 mRNA expression in peritoneal mouse cells following LPS injection ^13^. We and others demonstrated that Plk3 associates with calcium- and integrin-binding protein 1 (CIB1), and we showed that Plk3 kinase activity is regulated by CIB1 ^12, 14, 15^. Phosphorylation of Plk3 is regulated by protein phosphatase 6, though that phosphorylation does not appear to regulate its kinase activity ^11^.

In this report we reveal that Plk3 regulates the host response to enteric infection in mice. Mice deficient in Plk3 are highly resistant to models of systemic inflammation like CLP and CS injections that induce septic shock in WT mice. *Plk3^−/−^* mouse resistance is likely a result of limited bacteremia in the short term and the absence of bacteremia in the long term.

## Methods

### Reagents

All chemicals were purchased from ThermoFisher (Waltham, MA) unless otherwise stated. Anti-PRK antibody (Plk3) was purchased from Becton Dickinson (Franklin Lakes, NJ). JON/A antibody conjugated to PE was purchased from Emfret (Wurzburg, Germany).

### Mouse Strains

*Plk3^−/−^* mice were generated on a C57BL/6 background by Dr. Wei Dai at New York University and have been backcrossed 10 generations. WT control mice of the same background were purchased from The Jackson Laboratory (Bar Harbor, ME). Male and female mice aged 10-14 weeks were used for this study. All procedures were approved by the Thomas Jefferson University Institutional Animal Care and Use Committee.

### Hematologic Analysis

Whole blood from WT and *Plk3^−/−^* mice was collected via retroorbital bleed using a heparinized capillary tube. Blood cell analysis was carried out using a Hemavet blood cell analyzer (Drew Scientific, Miami Lakes, FL).

### Cecal Ligation and Puncture

Cecal ligation and puncture was performed according to Hubbard et al., with minor modifications ^16^. The cecum of age-matched WT and *Plk3^−/−^* mice were exposed and ligated just below the ileocecal valve. The cecum was punctured through and through with a 21-gauge needle and a small amount of cecal contents were extruded. The abdomen was closed in 2 layers and the mice were fluid resuscitated with 1 mL sterile saline. Mice were monitored and shock scores were recorded 24 hours post-surgery in a blinded fashion according to Mai et al. ^17^.

### Enzyme-linked immunosorbent assay

Plasma aspartate aminotransferase (AST) and alanine aminotransferase (ALT) were assessed using kits from Sigma-Aldrich (St. Louis, MO), and IL-1β was assessed using a kit from R&D systems (Minneapolis, MN).

### Cecal Slurry Injection

Cecal contents were isolated according to Star et al., with minor modifications ^18^. Ceca were isolated from WT donor mice, weighed, and diluted with water (500 μL/mg cecal contents). The resulting slurry was strained through an 860 μm mesh followed by a 190 μm mesh and mixed with an equal volume of 30% glycerol. Mice were injected IP with 10 μL/g body weight. Mice were monitored and shock scores were recorded in a blind fashion.

### Bacterial Load Assessment

To measure bacterial load in the peritoneum we first performed peritoneal lavage by injecting 5 mL ice cold saline into the peritoneum, modified from Walker W. E. ^19^. Fluid was then recovered, diluted in sterile PBS, and spread on blood agar plates. Plates were then incubated overnight at 37°C, and bacterial colonies were quantified. To measure bacterial load in the blood, retro orbital bleeds were performed, and blood was diluted 1:10 with sterile PBS. Diluted and whole blood were then spread on blood agar plates, incubated overnight at 37°C, and bacterial colonies were enumerated.

### Flow Cytometry

Flow cytometric assay of PE-JON/A binding to assess active α_IIb_β_3_ was performed using an Accuri C6 Plus (Becton Dickinson) as previously described ^20^.

### Lung Imaging

CLP was performed on WT and *Plk3^−/−^* mice following retro orbital injection (0.1μg/g mouse) of anti-mouse GPIX (Emfret) conjugated to AlexaFluor-750 (ThermoFisher) as previously described ^21^. 24 hours later the mouse was perfused with PBS and lungs were flushed with sterile PBS, fixed with paraformaldehyde, and inflated with agarose, then imaged using a LiCor (Lincoln, NE) Odyssey infrared imaging system. MFI was quantified using Odyssey software.

### Inflammatory cell quantification

Peritoneal lavage was conducted 6 hours after CLP and 100 μL were adhered to a microscope slide using a cytospin (Thermo). The cells were then stained with Diff-Quick (Siemens, Munich, Germany) and enumerated using an EVOS FL Auto microscope (Thermo).

### Statistical analysis

Statistical analyses were evaluated using Prism software (GraphPad). Survival curves were analyzed using a Mantel-Cox test while an unpaired Student’s T-test was used to evaluate the remaining data.

## Results

### Plk3 deletion confers protection from sepsis

It was reported that Plk3 is expressed in macrophages and that expression increases during attachment ^12^. In COS cells Plk3 expression is localized to the periphery ^12^. As macrophages are an essential component of innate immunity, we sought to determine if Plk3 regulates immune function. Therefore, we used *Plk3^−/−^* mice to assess the function of Plk3 in sepsis. We performed 100% cecal ligation and puncture (CLP) to produce severe polymicrobial sepsis and analyzed blood cell counts. We found that *Plk3^−/−^* mice were protected from leukopenia compared to WT controls, mostly because lymphocyte (LY) counts did not decrease to the same extent as those in WT mice as *Plk3^−/−^* white blood cell (WBC) counts were not statistically different from *Plk3^−/−^* sham blood cell counts, while WT WBC counts, and lymphocyte counts were greatly reduced compared to their sham counterparts (Table 1). As CLP results in enteric infection, we also analyzed inflammatory cell counts in the peritoneum of WT and *Plk3^−/−^* mice following sham or CLP surgery. We found that *Plk3^−/−^* mice had fewer WBCs in the peritoneum compared to WT mice following CLP, which was largely because fewer peritoneal neutrophils (NE) were present (Table 2).

**Table 1:** Blood cell counts in WT and *Plk3^−/−^* mice after CLP or sham surgery. * p < 0.05 *Plk3^−/−^* vs WT CLP, ^####^ p < 0.0001 WT Sham vs WT CLP, n = 7.

| Parameter | WT Sham | *Plk3*<sup>-/-</sup> Sham | WT CLP | *Plk3*<sup>-/-</sup> CLP |
| --- | --- | --- | --- | --- |
| WBC<br>k/μL | 3.71 ± 0.36 | 2.89 ± 0.38 | 0.70 ± 0.18 #### | 1.98 ± 0.61 * |
| NE<br>k/μL | 0.56 ± 0.22 | 0.85 ± 0.23 | 0.24 ± 0.13 | 0.49 ± 0.13 |
| LY<br>k/μL | 3.02 ± 0.26 | 1.97 ± 0.58 | 0.50 ± 0.12 #### | 1.30 ± 0.51 * |
| MO<br>k/μL | 0.12 ± 0.02 | 0.09 ± 0.05 | 0.10 ± 0.01 | 0.13 ± 0.07 |
| Plt<br>k/μL | 722 ± 65 | 627 ± 71 | 531 ± 70 | 534 ± 37 |

**Table 2:** Inflammatory cell counts in the peritoneum 24 hours following sham or CPL surgery. * p < 0.05 *Plk3^−/−^* vs WT CLP, ^#^ p < 0.05 vs corresponding sham value, ^##^ p < 0.01 vs corresponding sham value, n = 7.

| Parameter | WT Sham | *Plk3*<sup>-/-</sup> Sham | WT CLP | *Plk3*<sup>-/-</sup> CLP |
| --- | --- | --- | --- | --- |
| WBC<br>k/μL | 5.21 ± 0.63 | 3.23 ± 0.50 | 19.53 ± 5.65 <sup>##</sup> | 8.87 ± 1.05 *<br><sup>##</sup> |
| NE<br>k/μL | 2.31 ± 0.75 | 1.05 ± 0.24 | 12.3 ± 6.16 <sup>#</sup> | 5.27 ± 1.95 * <sup>#</sup> |
| LY<br>k/μL | 2.03 ± 0.76 | 1.65 ± 0.35 | 3.05 ± 1.85 | 2.02 ± 1.24 |
| MO<br>k/μL | 0.53 ± 0.08 | 0.30 ± 0.06 | 1.62 ± 0.80 | 1.01 ± 0.13 <sup>##</sup> |

To determine the consequence of these alterations in cell counts we performed CLP and monitored survival of WT and *Plk3^−/−^* mice. We found that all WT mice die by 48 hours following CLP, which is consistent with the literature ^22^. To our surprise, nearly all *Plk3^−/−^* mice survived over the course of the experiment (7 days) (Figure 1A). Plk3 deletion also protects mice from hypothermia following CLP (Figure 1B). Furthermore, 24 hours after surgery WT mouse shock scores indicate that all mice were in septic shock, while *Plk3^−/−^* mice were not in shock and had scores equal to that of sham operated mice (Figure 1C). We also found that the liver enzymes AST and ALT were greatly reduced in *Plk3^−/−^* mice 24 hours after CLP compared to WT mice, indicative of reduced liver damage (Figure 1D-E). Collectively, these data indicate that *Plk3^−/−^* mice are resistant to polymicrobial sepsis.

**Figure 1:**
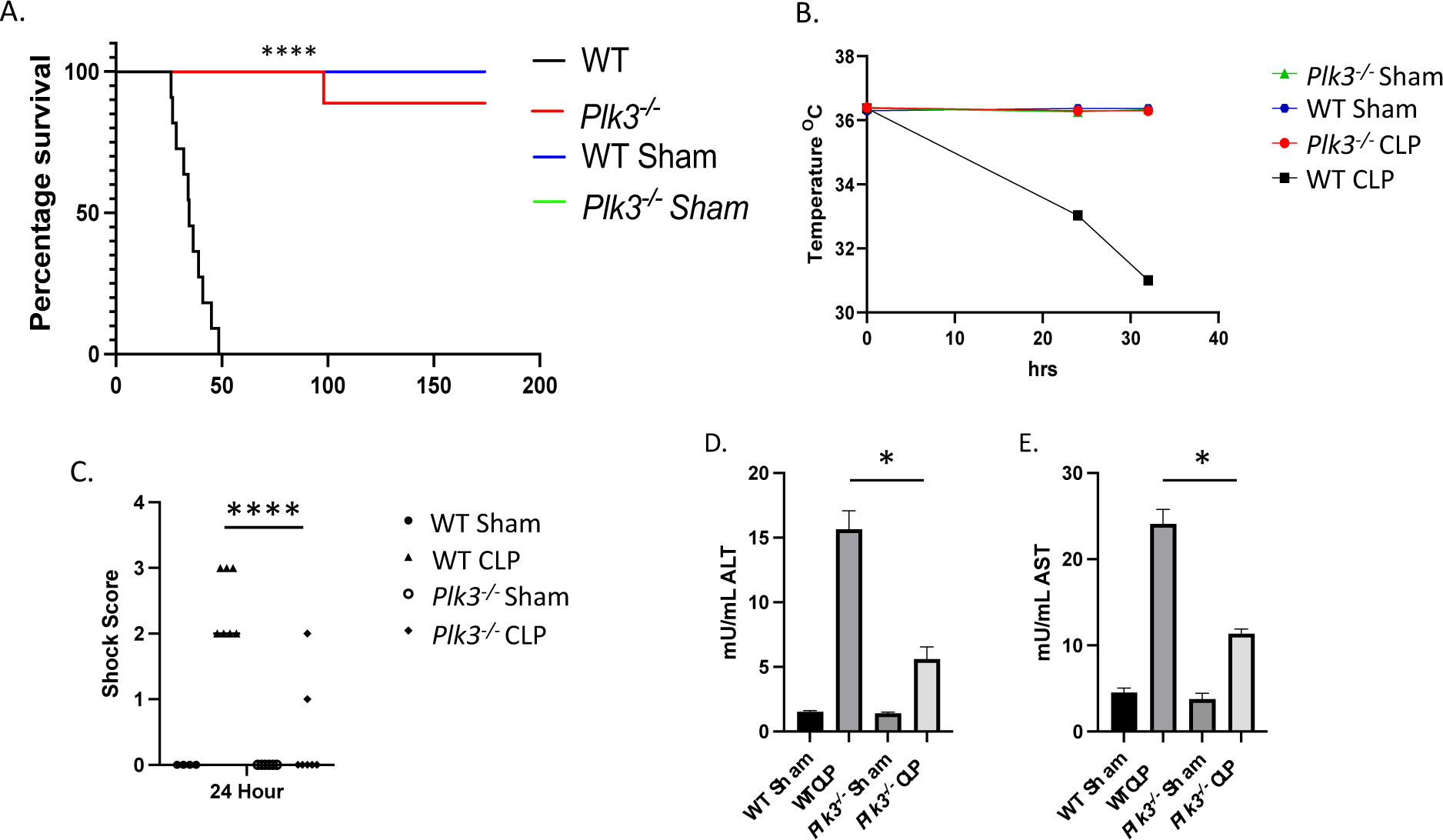
Plk3 deletion confers protection against sepsis. A) WT (n = 12) and *Plk3^−/−^* (n = 11) mice were subjected to severe sepsis via CLP and survival was monitored. B) Body temperature was recorded prior to CLP, 24 hours, and 32 hours after CLP. C) Shock scores were recorded 24 hours after CLP. D) Plasma alanine aminotransferase was assessed via ELISA 24 hours after CLP. E) Plasma aspartate aminotransferase was assessed 24 hours after CLP. **** p < 0.0001, * p < 0.05.

### Plk3^−/−^ mouse survival is independent of potential alterations to gut microbiota

It was reported that modifications of the mouse genome can influence gut microbiota ^23^. To determine if alterations in gut microbiota contribute to reduced mortality observed in *Plk3^−/−^* mice following CLP, we performed CS injections whereby cecal contents from WT mice were IP injected into *Plk3^−/−^* mice and vice versa. In each case our data mirrored that of CLP as recipient WT mice succumbed within 48 hours, while recipient *Plk3^−/−^* mice survived suggesting that changes to the gut microbiota are not responsible for *Plk3^−/−^* mouse survival following CLP (Figure 2A-B). Shock scores indicate that all WT mice injected with CS regardless of CS origin were in septic shock 24 hours after injection, while *Plk3^−/−^* mice were not (Figure 2C-D).

**Figure 2:**
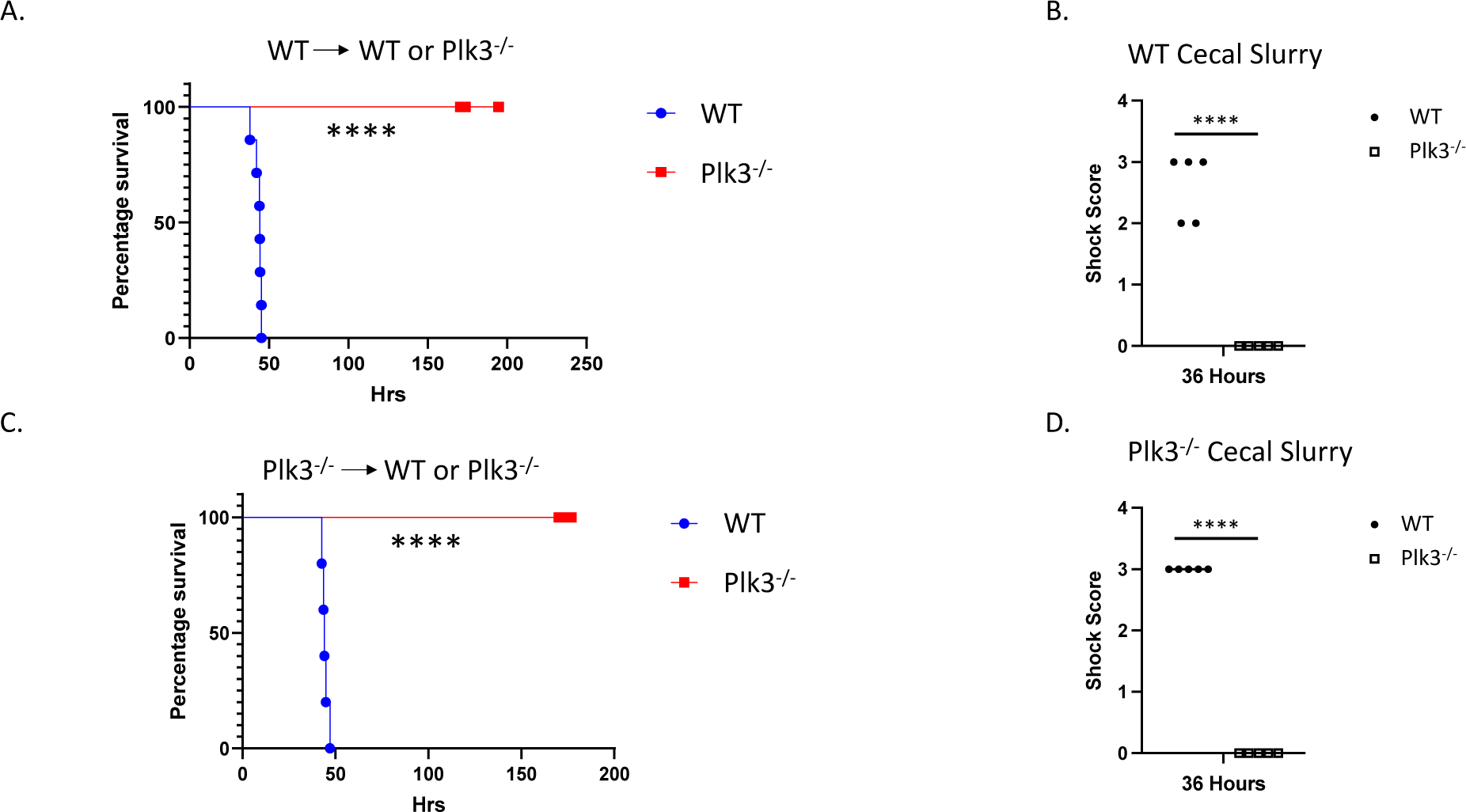
*Plk3^−/−^* mice survive CS injection. A) Cecal slurry was prepared from WT mice and injected into WT or *Plk3^−/−^* mice to evaluate survival (n = 7). B) Septic shock was analyzed 36 hours after injection. C) To rule out alterations in gut microbiota, cecal slurry was prepared from *Plk3^−/−^* mice and injected into either WT or *Plk3^−/−^* mice (n = 5). D) Shock scores were recorded 36 hours after injection. **** p < 0.0001.

### Plk3^−/−^ mice are resistant to systemic inflammation

Cytokine production is greatly enhanced during sepsis progression ^24^. Therefore, we examined plasma IL-1β in WT and *Plk3^−/−^* mice 18 hours after LPS injection. We found that mice lacking Plk3 had IL-1β levels comparable to that of vehicle (saline) injected mice, while WT mice had greatly increased IL-1β (Figure 3A). It is known that platelet reactivity is heightened during the initial phases of sepsis, which contributes to micro thrombus formation and eventually thrombocytopenia ^25, 26^. Engagement of TLR4, expressed on the platelet surface, results in immune responses as well as pro-thrombotic and pro-coagulation responses ^27^. Activation of platelet TLR4 leads to enhanced IL-1β expression ^28, 29^. Because IL-1β was quite low in *Plk3^−/−^* mice following LPS injection we determined if platelet activity was also altered following LPS (18 mg/kg) IP injection. We found that α_IIb_β_3_ activation, as measured by JON/A binding, was not enhanced in *Plk3^−/−^* mice injected with LPS, while it was greatly enhanced in WT controls (Figure 3A). To continue this line of investigation, we analyzed platelet deposition in the lung 24 hours after CLP in WT and *Plk3^−/−^* mice. We found significantly less platelet deposition in lungs from *Plk3^−/−^* mice compared to lungs from WT mice (Figure 3C-D). These data suggest that thrombosis is greatly reduced in *Plk3^−/−^* mice using a model of systemic inflammation and may help explain *Plk3^−/−^* mouse survival following CLP.

**Figure 3:**
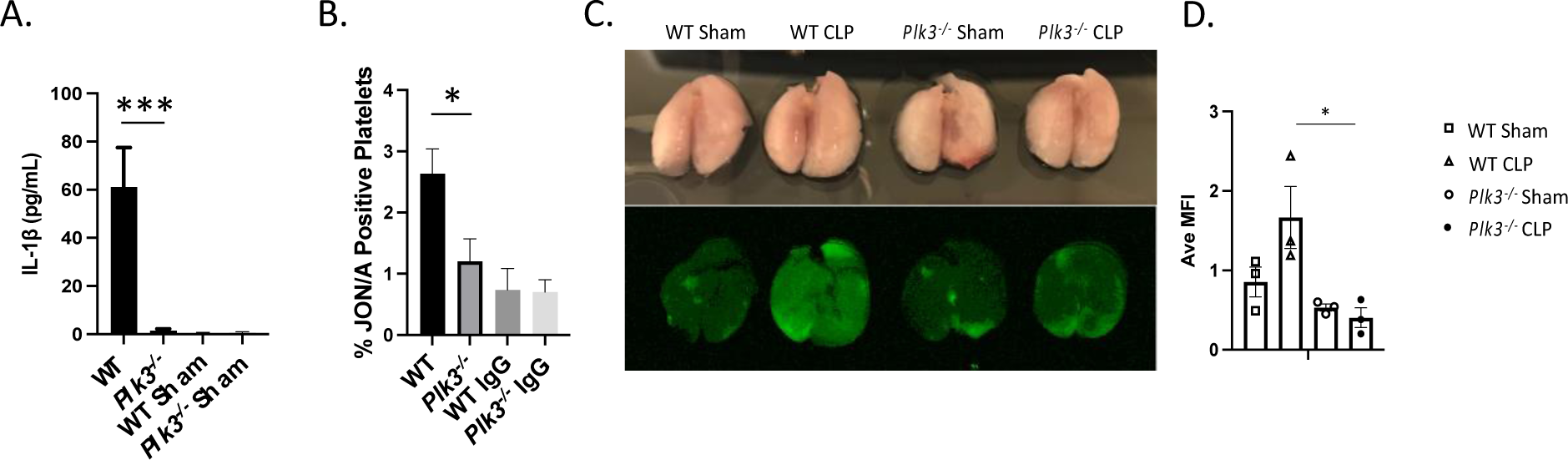
*Plk3^−/−^* mice are resistant to models of systemic inflammation. Mice were injected with 18 mg/kg LPS IP and blood was collected 18 hours later. A) Plasma IL-1β was assessed by ELISA (n = 3). B) JON/A binding was assessed via flow cytometry (n = 3). C) WT and *Plk3^−/−^* mice were injected with a fluorescent anti-GPIX antibody and lungs were excised 24 hours after CLP. D) Quantification of 3 independent experiments (n = 3). *p < 0.05, *** p < 0.001

### Reduced bacteremia with Plk3 deletion

For sepsis to occur bacteria must leave the site of the original infection and become systemic. To do that bacteria infiltrate the vasculature. Therefore, we assessed bacterial load following CLP in WT and *Plk3^−/−^* mice. Interestingly, an analysis of peritoneal bacterial load revealed that 24 hours after CLP, bacteria were numerous in the peritoneum of WT mice, while bacteria were virtually undetectable in the peritoneum of *Plk3^−/−^* mice (Figure 4A-B). Similar results were observed in the blood of WT and *Plk3^−/−^* mice (Figure 4C-D). These findings demonstrate that even over 24 hours very few bacteria are present in the blood in *Plk3^−/−^* mice. To determine if inflammatory cell migration is altered in *Plk3^−/−^* mice early after CLP, we quantified neutrophils and monocytes found in the peritoneum 6 hours post-surgery. We observed that neutrophil and monocyte migration was enhanced with Plk3 deletion and may explain the eventual clearance of bacteria following CLP (Figure 4E).

**Figure 4:**
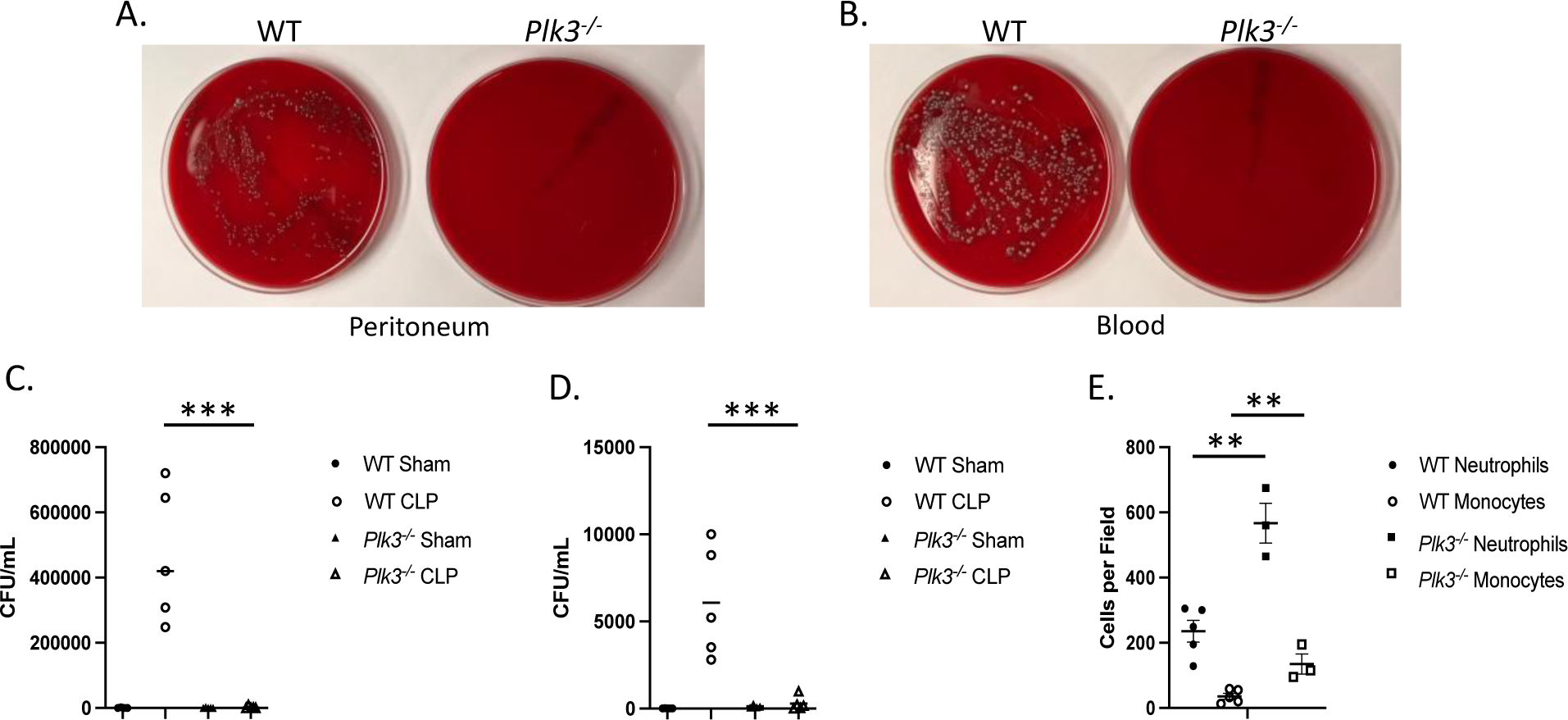
Bacteria are effectively cleared in *Plk3^−/−^* mice following CLP. A) Peritoneal lavage fluid was collected 24 hours after CLP and plated on blood agar and bacterial colonies were allowed to grow overnight at 37°C. B) Blood was drawn from WT and *Plk3^−/−^* mice 24 hours after CLP, plated on blood agar, and bacteria were allowed to grow overnight at 37°C. C) Bacterial CFU from the peritoneum of *Plk3^−/−^* and WT mice following CLP were quantified. D) Bacterial colonies from the blood of WT and *Plk3^−/−^* mice were quantified. E) Peritoneal lavage fluid was collected 6 hours after CLP, spun onto slides, stained, and cells were enumerated. **p < 0.01, ***p < 0.001.

To unravel this exciting finding, we tracked bacteria infiltration into the blood shortly after a lethal dose of CS, isolated from WT mice, that allows for tighter control over the enteric infection model. We found that a small number of bacteria were present in the blood of WT and *Plk3^−/−^* mice 60 minutes after injection with a trend towards an increase in WT blood (Figure 5A). However, bacteria infiltration in WT mouse blood increased greatly 180 minutes after injection but was not enhanced in *Plk3^−/−^* mouse blood (Figure 5B). Interestingly, the number of bacteria did not differ in the peritoneum of WT or *Plk3^−/−^* mice 60 or 180 minutes after injection (Figure 5A-B). These data suggest that even at early time points bacteremia is restricted in *Plk3^−/−^* mice even though plenty of bacteria are present in the peritoneum. This also suggests that the early migration of neutrophils and monocytes seen in *Plk3^−/−^* mice after CLP is not the reason for the limited bacteremia observed in these mice.

**Figure 5:**
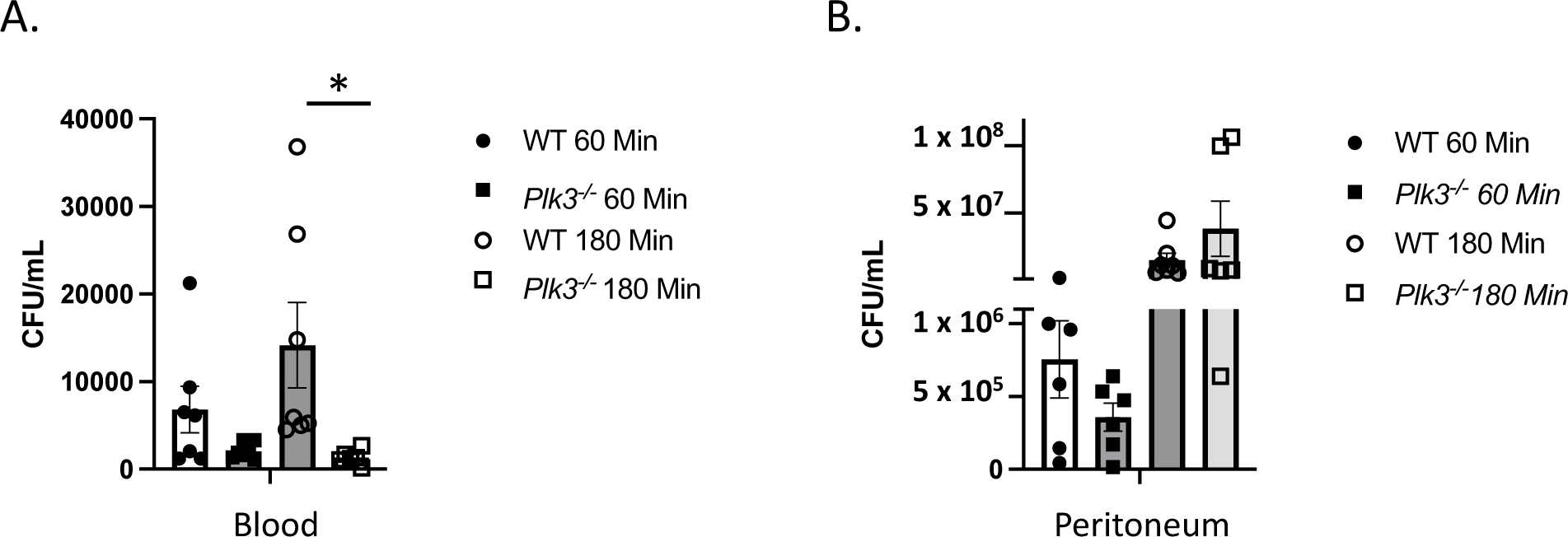
Bacteria fails to enter the blood of *Plk3^−/−^* mice following CS injection. A) Blood was collected from CS-injected WT and *Plk3^−/−^* mice 60 minutes and 180 minutes after injection, plated on blood agar overnight, and bacterial colonies were quantified. B) Similarly, peritoneal lavage fluid was collected from CS injected mice 60 minutes and 180 minutes post injection, plated, grown overnight, and quantified. * p < 0.05.

### Plk3^−/−^ mice are not protected from E. coli IV injection

To bypass the need for bacteria to infiltrate the blood from the peritoneum, we performed IV injection of GFP-*E. coli* into WT and *Plk3^−/−^* mice. Interestingly, Plk3 deletion conferred no protection against bacteria delivered in this manner and all *Plk3^−/−^* mice died along with their WT counterparts (Figure 6A). These data suggest that Plk3 dictates initial bacteria infiltration of the vasculature during infection and that protection from bacteria infiltration may be sufficient to prevent sepsis.

**Figure 6:**
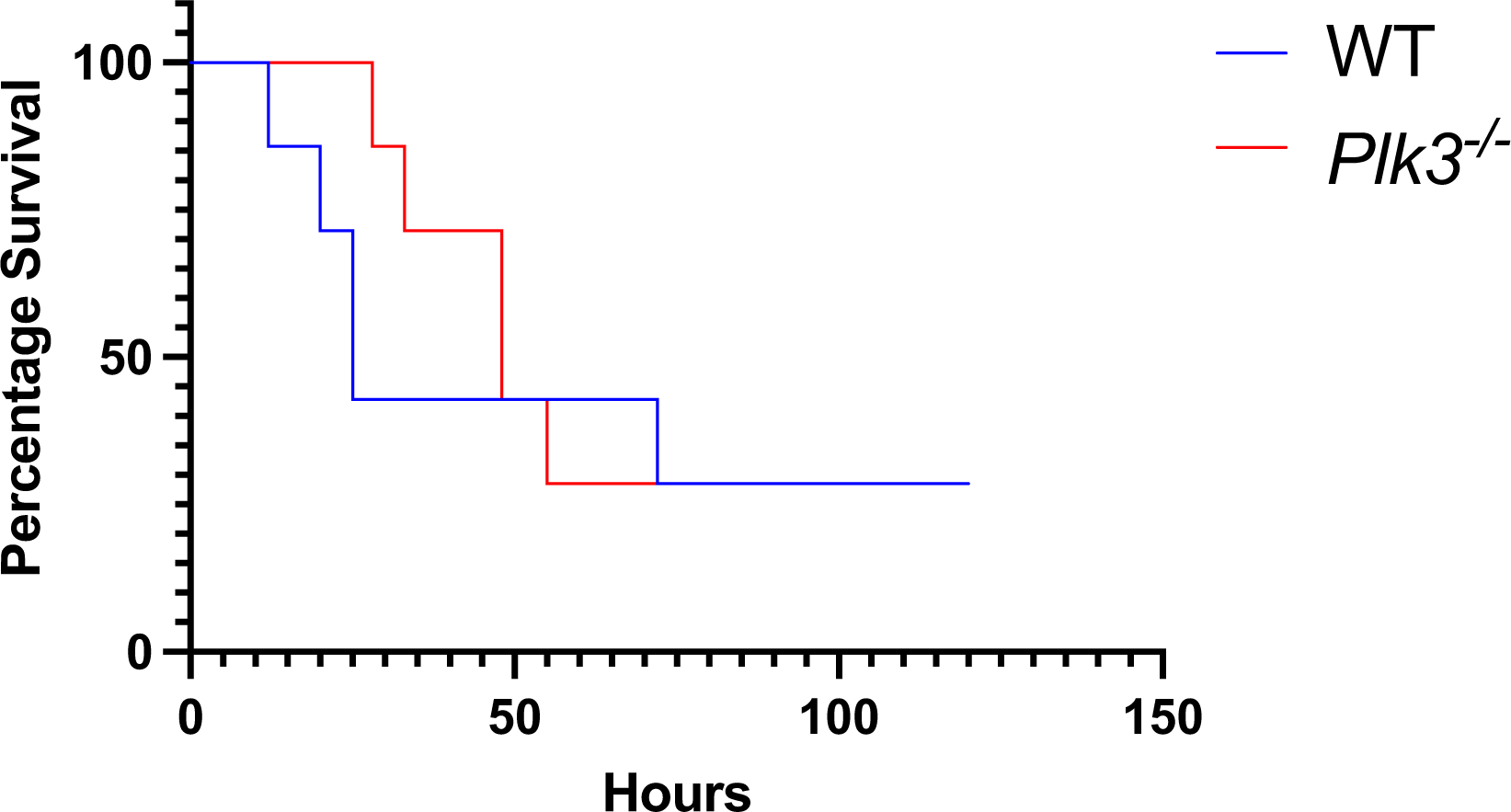
Plk3 deletion does not confer protection from I.V.-injected bacteria. WT and *Plk3^−/−^* mice were injected via retroorbital sinus with 1.5 X 10^8^ CFU *E. coli*. Survival was then monitored. n = 7, p = 0.6325.

## Discussion

In this report we demonstrate that Plk3 is a key regulator of bacteremia and that limiting Plk3 activity may improve outcomes for patients with infections that may otherwise lead to sepsis. We show that mice lacking Plk3 survive two models of severe sepsis (CS and CLP), while their WT counterparts do not. Our results suggest that this is because bacteria fail to invade the vasculature, an assertion that is supported by IV injection of bacteria, which results in death of *Plk3^−/−^* and WT mice. These data suggest that Plk3 regulates the mechanism by which bacteria enter the vasculature and that sepsis prevention may be possible by restricting bacteria to the original site of infection.

Sepsis can occur when a local infection becomes systemic, which means that bacteria must enter the vasculature and deposit in remote organs. Our data suggest that bacteria are restricted from moving from the peritoneum to the vasculature in *Plk3^−/−^* mice, even though there is no difference in the number of peritoneal bacteria between WT and *Plk3^−/−^* mice following CS injection. As *Plk3^−/−^* mice survival is not enhanced following IV injection of bacteria, the lack of bacteria in the blood following CS injection is likely not due to enhanced clearance, rather reduced vascular infiltration may be the cause. There are myriad examples in the literature that suggest vascular permeability is an important factor for sepsis progression. Reduction of vascular permeability during models of severe sepsis is protective ^30, 31^. Consistently, enhanced vascular permeability exacerbates sepsis ^32, 33^. In the case of *Plk3^−/−^* mice we would surmise that maintenance of endothelial integrity in the very early stages of enteric infection, prior to the development of sepsis, prevents dissemination of bacteria and results in nearly 100% survival after severe enteric infection.

We demonstrated that *Plk3^−/−^* mice maintain local control of enteric bacteria and resolve the local infection. Upon infection, inflammatory cells like neutrophils and monocytes respond quickly and are recruited to protect the host ^34–36^. It is generally accepted that neutrophils are recruited to the site of infection first, followed by the arrival of monocytes. Proper neutrophil function is critical to avoid sepsis, and impairment of neutrophil function is often observed in cases of severe sepsis in animal models ^37, 38^. In the case of *Plk3^−/−^* mice however, peritoneal bacteria numbers three hours after CS injection are not reduced compared to WT control mice suggesting that enhanced phagocytosis in the peritoneum is not responsible for the lack of bacteria found in the blood of *Plk3^−/−^* mice in early time points after CS injection.

Neutrophils and monocytes begin to arrive in the peritoneum six hours after CLP surgery. At that time, we noticed that more of each cell type were present in *Plk3^−/−^* peritonea than WT peritonea. It is likely that this enhanced migration of inflammatory cells is responsible for the eventual resolution of the infection 24 hours post-surgery. Reduced bacteremia, as observed in *Plk3^−/−^* mice following enteric infection, may protect the host from the robust inflammatory response associated with the progression of sepsis, ensure proper function of inflammatory cells, and allow the host the resources to combat the local infection. In fact, 24 hours after surgery there are fewer inflammatory cells present in *Plk3^−/−^* peritonea than WT peritonea because the infection has been resolved in *Plk3^−/−^* mice while it persists in WT mice. Consistently, *Plk3^−/−^* mice are protected from leukopenia likely because the inflammatory response associated with sepsis is absent.

Plk3 has been connected to inflammatory cell function as Plk3 mRNA and protein expression is enhanced in human macrophages as they transition from peripheral monocytes to attached macrophages ^12^. Similarly, Plk3 mRNA expression is quickly elevated in LPS-treated mouse peritoneal cells ^13^. These data suggest that Plk3 expression is enhanced during infection or the early stages of sepsis in both mouse and human. A comparison of mouse and human cells treated with LPS, suggests that the transcriptional response is very similar in mouse and human during systemic inflammation caused by LPS ^13^. These data highlight the similarity of the response to infection in mouse and human and suggest that Plk3 may be a viable biomarker for sepsis in patients.

In conclusion, we show that Plk3 is an important regulator of bacteremia and that deletion of Plk3 limits bacteremia in the host to a degree that allows the host to respond to the original infection. Deletion of Plk3 confers nearly complete protection from polymicrobial sepsis caused by enteric infection. Our data point to Plk3 as a promising therapeutic target for the treatment of sepsis.

## Acknowledgments

We would like to acknowledge Antonios Tawk, MD for his work on the IL-1β ELISA

## Non-standard Abbreviations and Acronyms

Plk3: polo-like kinase 3
WT: wildtype
CLP: cecal ligation and puncture
CS: cecal slurry
PAMP: pathogen associated molecular pattern
LPS: lipopolysaccharide
PRR: pattern recognition receptor
TLR4: toll-like receptor 4
GFP: green fluorescent protein
IL: interleukin
CIB1: calcium- and integrin-binding protein 1
PE: phycoerythrin
AST: aspartate aminotransferase
ALT: alanine aminotransferase
PBS: phosphate buffered saline
GPIX: glycoprotein IX
WBC: white blood cell
LY: lymphocyte
NE: neutrophil
MO: monocyte
Plt: platelet

